# A bio-informatics approach to identify new drug targets in multidrug-resistant bacteria

**DOI:** 10.1101/2025.05.31.657076

**Authors:** Imogen Bramhill, Aryeh Chiam, Dominique de Jong-Hoogland, Martin B. Ulmschneider

## Abstract

Antibiotic resistance poses a global health crisis. In order to develop new antibiotic agents, it is crucial to identify drug targets in multidrug-resistant bacteria. Criteria for such a target are an α-helical, essential membrane protein, that is non-homologues with the human membrane proteome, and present across multiple bacterial species. Using a stepwise subtractive genomics approach, the membrane protein F0F1 ATP synthase subunit C was identified as a non-human analogues drug target that is present in 11 bacterial species.

## Introduction

Antibiotic resistance is the root cause of a rapidly worsening global public health crisis. In 2019, Antimicrobial Resistance Collaborators found that drug-resistance infections killed at least 1.27 million people worldwide, and nearly 5 million deaths associated with bacterial antimicrobial resistance (AMR).^1^ With predictions raising the number of deaths due to bacterial AMR to 10 million deaths annually by 2050,^2^ the UN Environment Programme, the US Centers for Disease Control and Prevention (CDC), World Health Organisation (WHO) and European Commission have recognised the need for urgent global action.^3-7^

The cost of antibiotic research and low rate of novel antibiotic translation chiefly due to the extremely unfavourable cost-benefit ratio, has discouraged big pharmaceutical companies from developing new antibiotics.^8-10^ Consequently, only 17 new antibiotics have been approved between 2010 and 2021,^1, 11^ five of which are not in circulation due to the companies’ finances.^8, 9^ Only four of these new antibiotics represent new classes of antibiotics, targeting bacteria through novel mechanisms.^12, 13^ Though there have been three new FDA-approved antibiotics last year (2024), experts worry that this could be a peak due to their associated scientific, regulatory and financial challenges.^14^ It remains that many of the broad-spectrum antibiotics currently in use are not effective against multidrug-resistant bacteria, thus it is critical to look for novel drug targets specific to these species. In 2017, the WHO published a priority pathogens list for research and development of new antibiotics^5^, updated in 2024,^15^ and dominated by Gram-negative bacteria. This list was composed of multidrug-resistant bacteria – against some of which even the last-resort antibiotic colistin is ineffective.^16, 17^ This list was used as the starting point of this investigation which endeavours to identify new drug targets to combat multidrug-resistant bacterial infections.

Membrane proteins constitute the largest class of drug targets as they are responsible for vital physiological processes such as nutrient and waste transportation, signal transduction, and enzymatic catalysis.^18, 19^ Due to their placement in the membrane, membrane proteins fold in a very specific way so that the hydrophobic residues are buried in the hydrophobic membrane core, while the more polar residues tend to reside either side of the membrane where they are exposed to aqueous environments.^20^ Membrane proteins divide overwhelmingly into two major structural classes, which reflects the two secondary structural motifs that are stable in lipid bilayer membranes: the alpha (α)-helix and the beta (β)-barrel.^21, 22^ α-helical proteins are found in the inner membranes of Gram-negative bacteria, Gram-positive bacteria, and eukaryotes while β-barrel proteins are almost exclusively found in the outer membranes of Gram-negative bacteria.^23-25^ In an effort to identify a new target for broad-spectrum antibiotics this investigation will focus on α-helical transmembrane proteins.

## Methods

A stepwise subtractive genomics approach was used for the identification of unique essential membrane proteins in the 21 bacterial species selected based on the 2017 WHO priority pathogens list, including 8 species of the Enterobacteriaceae family and subspecies of Shigella and Providencia.^5^ The scientific names and accession numbers are listed in **Table 1**. Proteins were clustered according to sequence similarity across species to identify potential drug targets in multiple bacterial species, and the protein sequences were aligned and analysed. The overall workflow is illustrated in **Figure 1**.

**Table 1.** The chosen 21 species of bacteria with accession numbers and (-) or (+) indicating Gram-negative or Gram-positive, respectively. Based on 2017 WHO priority pathogens list and ordered accordingly.

| Species | Accession Number | Group |
| --- | --- | --- |
| *Acinetobacter baumannii* | NZ_CP015121.1 | Bacteria(-) |
| *Pseudomonas aeruginosa* | NC_002516.2 | Bacteria(-) |
| *Enterobacter cloacae* | NZ_CP009756.1 | Bacteria(-) |
| *Escherichia coli* O157 | NC_002695.2 | Bacteria(-) |
| *Klebsiella pneumoniae* | NC_016845.1 | Bacteria(-) |
| *Morganella morganii* | NZ_CP034944.1 | Bacteria(-) |
| *Proteus mirabilis* | NC_010554.1 | Bacteria(-) |
| *Providencia rettgeri* | NZ_CP029736.1 | Bacteria(-) |
| *Providencia stuartii* | NC_017731.1 | Bacteria(-) |
| *Serratia marcescens* | NZ_CP063354.1 | Bacteria(-) |
| *Enterococcus faecium* | NZ_CP039729.1 | Bacteria(+) |
| *Staphylococcus aureus* | NC_007795.1 | Bacteria(+) |
| *Helicobacter pylori* | NC_017379.1 | Bacteria(-) |
| *Campylobacter jejuni* | NC_002163.1 | Bacteria(-) |
| *Salmonella typhimurium* | NC_003197.2 | Bacteria(-) |
| *Neisseria gonorrhoeae* | NZ_CP012028.1 | Bacteria(-) |
| *Streptococcus pneumoniae* | NZ_CP007593.1 | Bacteria(+) |
| *Haemophilus influenzae* | NZ_CP009610.1 | Bacteria(-) |
| *Shigella flexneri* | NC_004337.2 | Bacteria(-) |
| *Shigella dysenteriae* | NZ_CP061527.1 | Bacteria(-) |
| *Mycobacterium tuberculosis* | NC_000962.3 | Bacteria(-) |

**Figure 1.**
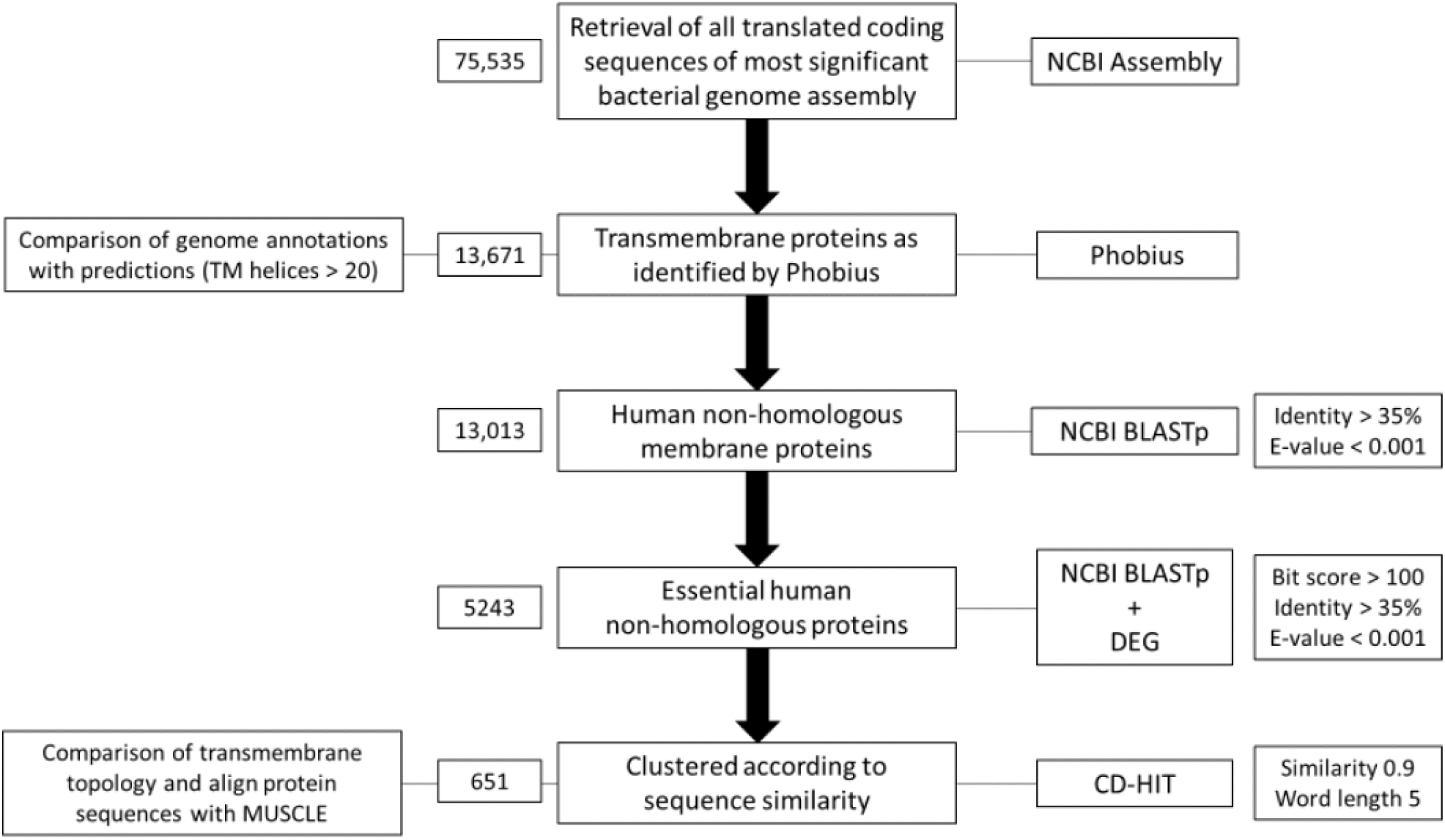
Overall workflow chart. Bit score is a measure of sequence similarity independent of query and database size (the higher the better for this study); E-value is the number of expected hits whih would be found randomly (the lower the better for this study).

### Retrieving bacterial genomic data and human proteome

The most significant complete genomes (e.g. reference or representative genomes) of the chosen 21 bacterial species were retrieved as translated coding sequences in FASTA files from the NCBI Assembly database at https://www.ncbi.nlm.nih.gov/.^26^ The human reference genome GRCh38 was retrieved in the same format.^27^

### Identification of membrane protein sequences

The translated coding sequences of each bacterial species and the *Homo sapiens* GRCh38 reference genome were analysed using Phobius algorithm at https://phobius.sbc.su.se/, to locate and predict the topology of trans-membrane (TM) segments, thus identifying membrane proteins^28^. In assembly entries with chromosomal and plasmid DNA, the plasmid sequences were ignored due to their transiency and inconsistency between strains. The proteins predicted to contain TM segments were taken forward for further analysis, though sequences with predicted signal peptides were not taken further due to possible inaccurate predictions.

A Python script was written to create a **dictionary** of the translated coding sequences and their genomic annotations for the human and bacteria proteomes, which created a new FASTA file containing only the membrane protein sequences. These FASTA files were used in the next stages of comparison genome analysis using local BLASTp^29, 30^.

### Identification of non-homologous proteins

A local NCBI BLASTp searching for non-homologous bacterial membrane proteins was conducted with E-value cut-off of 0.001 and 35% identity^31-36^ against the human membrane proteome.^29, 30, 37^ The homologous proteins were excluded, and the non-homologs merged into a list, by parsing the outputs in formats 0 and 6.

### Identification of essential non-homologous proteins

The human non-homologous membrane proteins were subjected to a local BLASTp similarity search – with E-value cut-off of 0.001, 35% identity and bit-score cut-off of 100 – against the Database of Essential Genes (DEG) to focus on the proteins required for the survival of the bacteria.^38^ The parameters used were similar to filtering out the human homologous proteins, with the additional bit-score cut-off achieving closer matches to identify essential proteins. This greater similarity threshold has been used by previous computational studies identifying potential drug targets in bacteria.^31-36, 39-42^

### Clustering of membrane proteins

The essential non-homologous membrane proteins were subjected to the CD-HIT suite with a cut-off score 0.9 (90% sequence identity) to cluster the proteins across bacterial species according to sequence similarity.^43-46^ Clusters containing ≥5 proteins were selected for sequence analysis; the protein ID headers were paired with the protein sequences and genome annotations used to record the protein names for each cluster.

### Sequence alignment and analysis of TM segments

The consistency of the genome annotations of proteins in clusters of ≥5 proteins indicates function similarity, so the proteins used for sequence alignment and analysis were those with consistent genome annotations across species. The largest protein clusters were prioritised with each protein’s sequence in the clusters being aligned using MUSCLE^47^. The multiple sequence alignments, saved as CLUSTALW (.aln) files, were viewed using UGENE^48^ and predicted topologies from Phobius were used as a guide to compare sequences and infer conserved structure.

### Metabolic pathway analysis

The representative protein sequences of each of the 109 protein clusters were concatenated into one FASTA file and run through the BLASTKOALA server^49^. The genes were functionally characterised by BLASTp and KEGG orthology mapping^50-52^ to identify the pathways for the proteins showing their most likely role in the organism.

### 3D protein structures (PyMOL)

The structures containing proteins SecY and Sec61 were downloaded as PDB files from the RCSB PDB website (rcsb.org)^53, 54^ with the PDB IDs 3DIN^55, 56^ and 6R7Q^57, 58^. The structures were viewed with PyMOL^59^ and the chains of SecY and Sec61α were extracted to separate objects from the complexes in 3DIN and 6R7Q, respectively: Chains B/C and F for SecY in 3DIN and Chain HC/XX for Sec61α in 6R7Q. The two identical chains of SecY were superimposed to show the similarity, and SecY and Sec61α aligned in PyMOL to compare experimental protein structure.

## Results and Discussion

All 75,535 translated coding sequences from the chosen bacterial genomes were filtered down after running through Phobius to leave just 13,671 membrane proteins’ sequences that did not include signal peptides. The percentage of the genomes encoding membrane proteins was calculated for each bacterium and averaged across all species, resulting in 18% which closely matches the 19% of transmembrane proteins in the human proteome. Though TM proteins containing signal peptides (SPs) were included in this statistic, they were not taken forward for further analysis as SPs may be cleaved or form part of an excreted protein; the number of TM proteins with a signal-peptide compared to the total number of TM proteins in each species of bacteria is shown in **Table S1**.

The set of TM proteins was analysed to determine the number and distribution of TM helices. The results of this quantification were summarised in **Figure 2**. Proteins found to be single-spanning (i.e., only one α-helical segment traversing the membrane) were not taken forward as the simpler structure tends to provide fewer options for a drug binding site.^60, 61^

**Figure 2.**
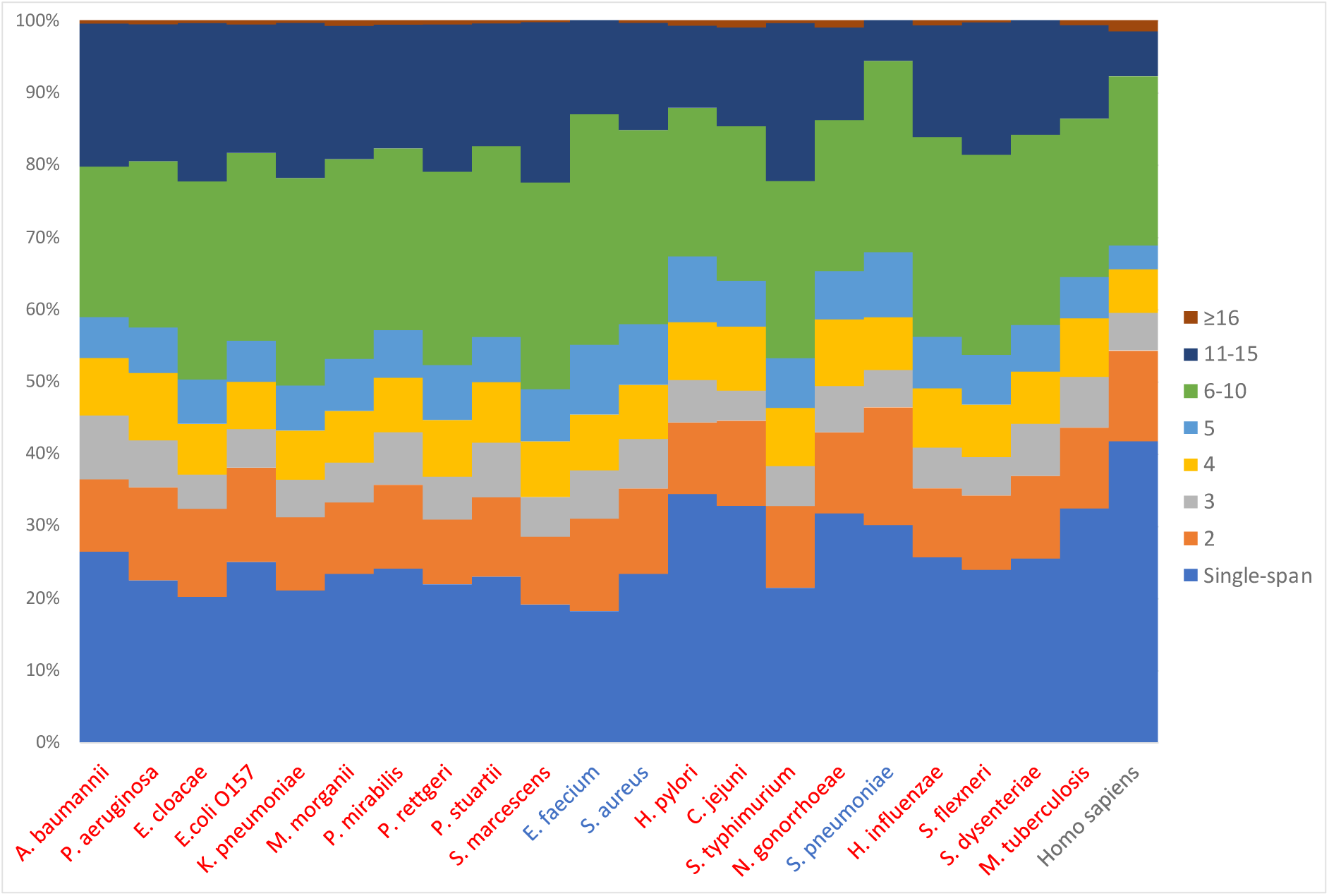
Transmembrane helices distribution in bacterial and human membrane proteins. Bar charts showing the distributions of helices among membrane proteomes in each species of bacteria (gram-negative labelled in red, gram-positive in blue) and in humans (*Homo sapiens* – labelled in black).

Analysis of the distribution of the number of TM helices among membrane proteins was repeated on the human proteome after extracting them from Phobius results. Like the distributions found in bacteria, single-spanning proteins were the most dominant by far, rising to 42% of the whole membrane proteome (see **Figure 3**).

**Figure 3.**
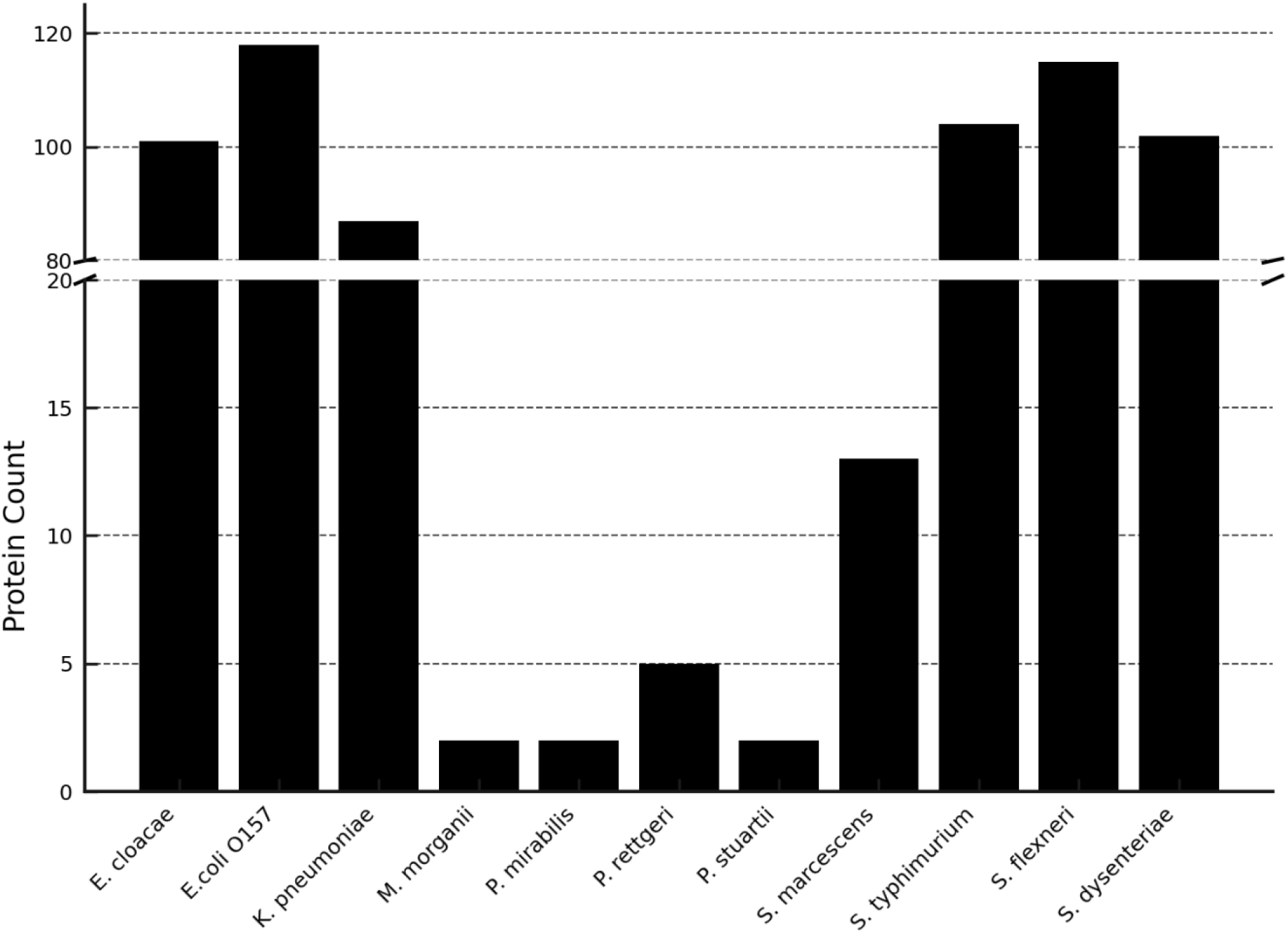
Bar chart showing the populations of clusters containing ≥5 proteins with ≥ 90 similarity across different species of bacteria, bacteria absent from the clusters are omitted.

The local BLASTp searches conducted with each bacterial membrane protein FASTA file against the human membrane proteome reduced the number of sequences by around 5%. The thresholds used here were informed from previous work reporting subtractive genomic analysis of other bacteria.^31, 35, 41^ The sets of non-homologous proteins included proteins labelled as “No hits” as well as those which satisfied the thresholds. The BLASTp tool was used for a selective approach with the 21,967 membrane protein sequences being compared like for like.

The membrane proteins found to be non-homologous with the human membrane proteome, were subjected to a local BLASTp search against the Database of Essential Genes (DEG) which utilises the minimal-gene-set concept for bacteria^38, 62^; this process revealed about 40% of them to be genes required to sustain life. The proportion of essential to non-essential membrane proteins is shown in **Figure S1**. These remaining proteins are both non-homologous with the host proteome and essential for survival of the bacteria – 2 key aspects which make for species-specific potential drug targets.

Having gathered a set of human non-homologous essential proteins in each of the chosen 21 bacterial species, the aim was to find a drug target that is present across species. To narrow down the number of proteins for this purpose, they were clustered according to sequence similarity using CD-HIT suite.^43, 44^ A threshold of 80% sequence similarity was initially used since membrane proteins are known to have highly conserved structures due to their restricted folding in the membrane, despite having distinct functions in the cell; therefore, this higher than typical threshold ensured proteins with both similar structure and function were grouped together, while reducing noise.^21, 63, 64^ However, latterly the need for sequence alignment required a threshold of 90% similarity. To extract the full protein ID of every protein in the clusters and for retrieval of the protein sequences for sequence analysis later, additional settings were used – the details can be found on Github.^43^ Protein sequences containing non-letter characters (i.e. sequencing gaps) were removed, leaving 5183 proteins for clustering.

There were 109 clusters which contained 5 or more proteins, the clusters were split into their own files of sequences and accession numbers for sequence analysis and comparisons. The output file from the CD-HIT clustering can be found as a CSV file in the linked data repository (**Dataset D1**). Collating the data showed the distribution of bacterial species in these clusters (see **Figure 3**) where 6 Gram-negative bacteria appeared the most frequently: *E. cloacae, E. coli, K. pneumoniae, S. typhimurium, S. flexneri* and *S. dysenteriae*. There were 13 clusters which contained these 6 bacteria in addition to *Serratia marcescens*; *M. tuberculosis* did not share similar membrane proteins with other bacteria. There were 2 protein clusters containing 11 bacterial species, suggested from genome annotations to be preprotein translocase SecY and F0F1 ATP synthase subunit C, which were investigated latterly as potential drug targets.

The clustered protein sequences aligned by MUSCLE were visualised in UGENE, to infer the structural similarity among each cluster – the alignment files can be found in the linked data repository (**Dataset D2**). This high conservation among many of the TM sequences further supports the homology of the proteins in each cluster. As sequence leads to structure, which encodes function, these unique essential proteins are potential novel antibiotic targets, a hypothesis that is supported by the functions listed in the genome annotation.

After completing the analysis, Kyoto Encyclopaedia of Genes and Genomes (KEGG) was used to look into the metabolic pathways of the 109 sets of proteins found.^32, 33, 41^ The KEGG Pathway Analysis results overview broken down by category and subcategories are summarised in **Table S3**. The 109 representative sequences were input as the query and 8 were eliminated due to the absence of recognised metabolic pathways (see **Table S2**), leaving 101 possible proteins across 5 or more high-priority antibiotic-resistant bacterial species. A summary of the results of using BLASTKOALA^49^ are reported in **Table 2**, showing the distribution of metabolic pathways. The most highly populated functional categories were signalling and cellular processes and environmental information processing with 29 and 28 proteins, respectively. Proteins were involved with biochemical pathways such as oxidative phosphorylation and membrane transport, with 26 ABC transporters reported.

**Table 2.** Summary of the results of the metabolic pathway analysis through BLASTKOALA showing the number of proteins in each functional category.

| BLASTKOALA Category | Number of Proteins |
| --- | --- |
| Protein families: signalling and cellular processes | 29 |
| Environmental Information Processing | 28 |
| Energy metabolism | 10 |
| Carbohydrate metabolism | 8 |
| Protein families: genetic information processing | 5 |
| Genetic Information Processing | 5 |
| Glycan biosynthesis and metabolism | 4 |
| Unclassified: signalling and cellular processes | 3 |
| Protein families: metabolism | 3 |
| Unclassified | 2 |
| Unclassified: metabolism | 2 |
| Human Diseases | 1 |
| Lipid metabolism | 1 |

The protein clusters were evaluated for conservation among bacterial species as ideally a drug could be used to target more than one species of antibiotic-resistant bacteria. Starting with proteins conserved in 5 species, namely *E. cloacae, E. coli, S. typhimurium, S. flexneri* and *S. dysenteriae*, there were 36 proteins identified as potential membrane protein drug targets (see **Table 3**). Looking at non-homologous membrane proteins essential and common to 6 species – these same 5 species and *K. pneumoniae* – there were 47 potential antibiotic targets identified (see **Table 3**). While these 6 species appeared most often in the protein clusters, there were 10 proteins found in these species that were also present in *S. marcescens* (see **Table 3**). These are dominated by permease and other transport proteins which would be a significant target for antibiotics as inhibition would prevent vital biomolecules and ions from reaching the cell.

**Table 3.**
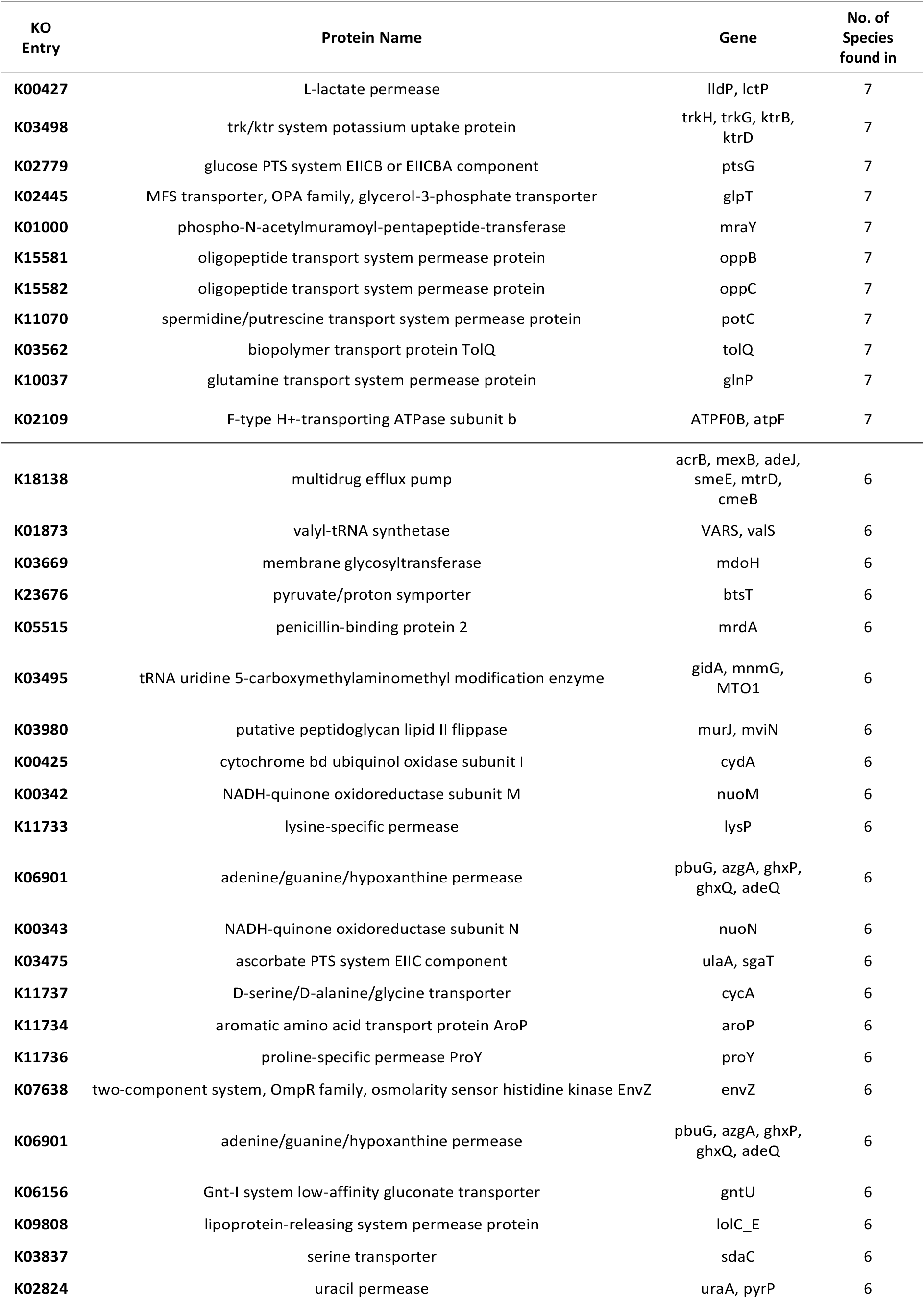
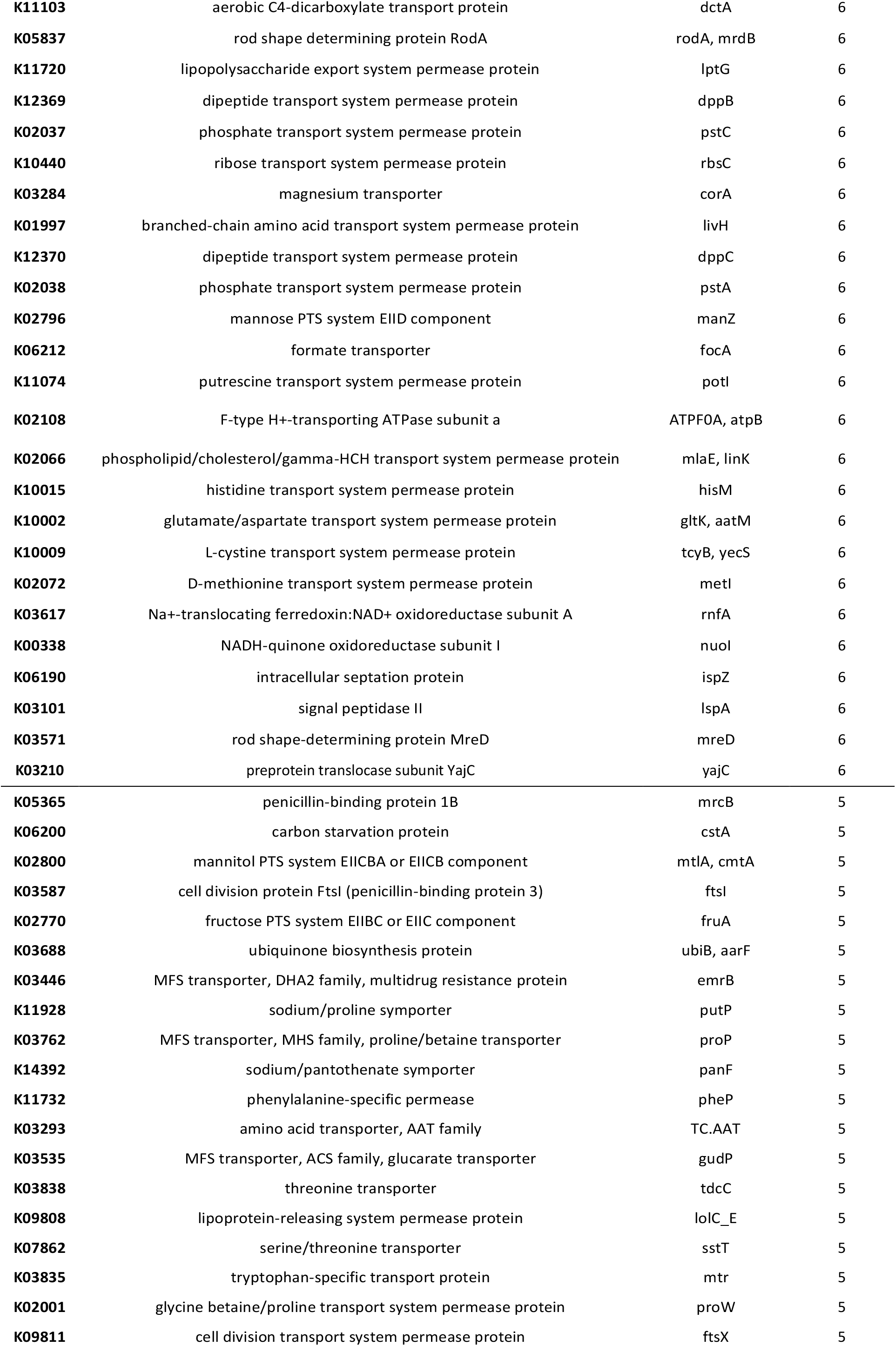
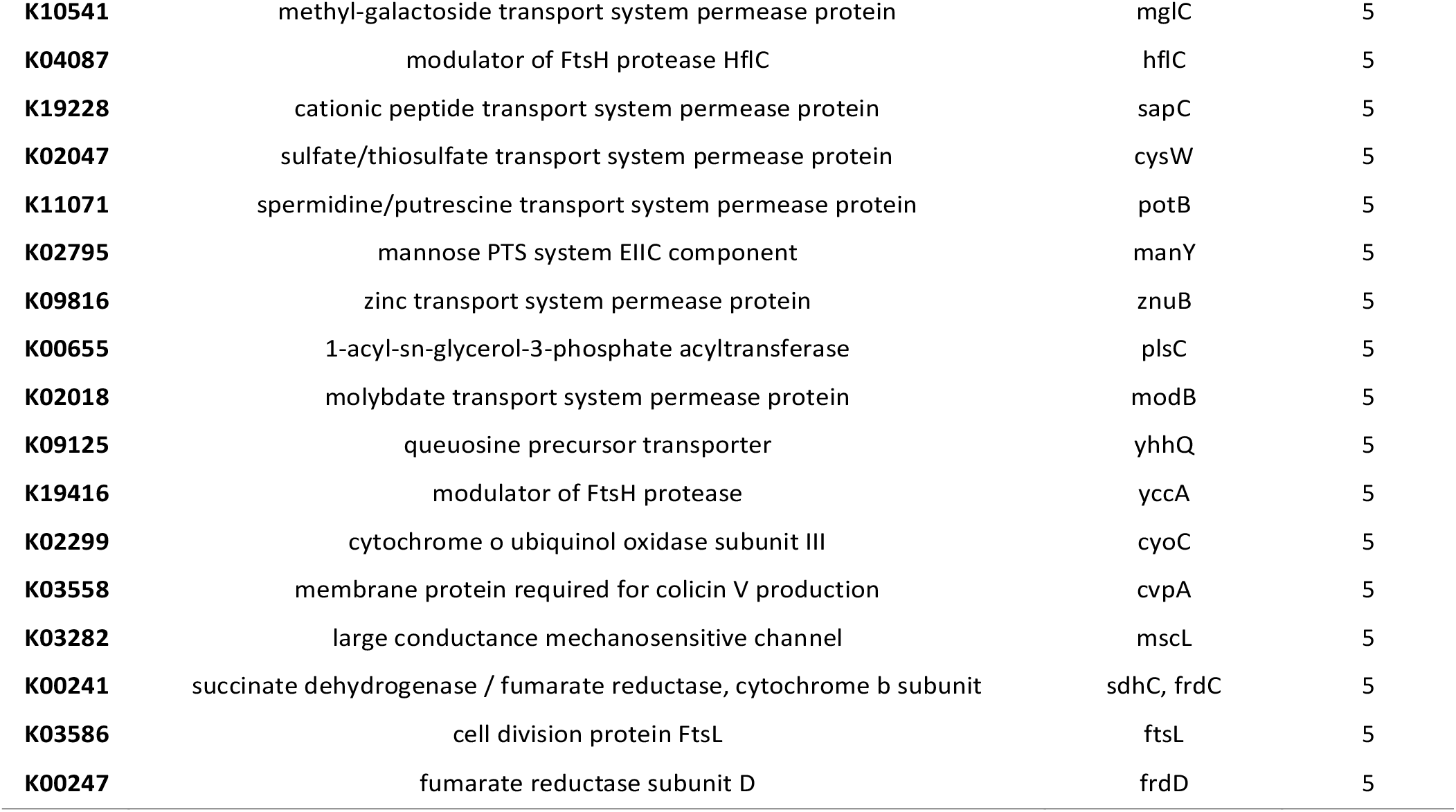
Protein names, K number and gene from KEGG orthology annotation for proteins found in 5-7 species.

Although there were a potential 109 proteins to work with as an output from the protein clustering, some candidates were eliminated where annotated gene functions differed across species or metabolic pathways could not be obtained, justification for elimination shown in **Table S2**.

There were only 2 membrane proteins found to be common to 11 species, following the workflow of this study: SecY and F0F1 ATP synthase subunit C. SecY has been cited in literature as a homolog of the human translocase subunit Sec61α,^65^ so the translated coding sequences for Sec61α were extracted from the GRCh38 genome and aligned with the SecY protein sequences. The overall percentage sequence similarity between S. flexneri protein NP_709088.1 and the 4 Sec61α isoforms was calculated in UGENE, with protein NP_709088.1 chosen as the reference sequence from being set as the representative sequence of the cluster by the CD-HIT suite. Across the 4 isoforms of human protein Sec61 subunit α, the similarity was between 14-16% excluding gaps, supporting the result of the initial BLASTp search against the human proteome. However, characterised structures of SecY and Sec61α were obtained from the RCSB PDB database.^53-58^ The SecY pdb file came as a dimer so one chain was extracted for structural comparison; the SecY and Sec61α structures were superimposed (see **Figure 4**). Although not identical, it is clear there are many general structural similarities: though the cytoplasmic and extracellular loops differ – e.g. the intracellular loop at residues 231 to 271 in SecY compared to Sec61α loop (G260 to P290), the TM helices share a very similar structure. This visual comparison could point to the extracellular loops at the top of the proteins shown in **Figure 4** being a possible unique binding site for a drug due to their differences. Further comparative analysis of these 2 proteins should be carried out to determine if SecY remains a potential non-homologous drug target for 11 of the priority pathogens studied here. This inconsistency of sequence similarity and structural similarities may highlight the limitations of purely sequence-based analyses.

**Figure 4.**
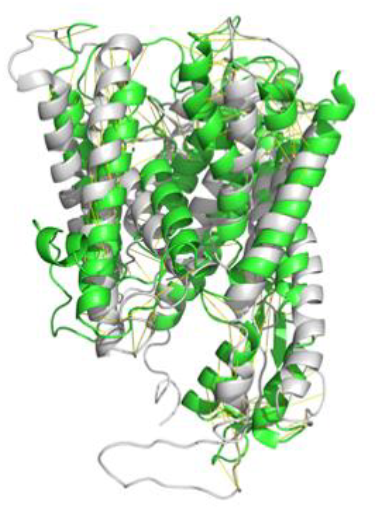
Tertiary structure overlay of SecY in gray (pdb: 3DIN) and Sec61α in green (pdb: 6R7Q).

F0F1 ATP synthase subunit C, though only 79 amino acids long, has a highly conserved structure, possibly a dimeric structure with similar patterns of glycine and alanine residues in both TM segments (see **Figure S2**). According to the KEGG orthology analysis, this protein is classified under “Metabolism”, associated with essential pathways such as oxidative phosphorylation (energy generation) and photosynthesis (light to chemical energy conversion). Since there is already a class of clinically used antibiotics, diarylquinolines, that targets the ATP synthase subunit C in *M. tuberculosis*, the F0F1 ATP synthase subunit C found across the 11 species is a potential drug target.

## Conclusion

In conclusion, being human non-homologous and present in 11 of the 21 investigated priority bacteria, F0F1 ATP synthase subunit C may be a target for potential therapeutic drug development in future. In addition, this study yielded 99 potential membrane protein targets for antibiotics against 5-7 species of multidrug-resistant bacteria. However, homology modelling, binding site analysis and *in vivo* experimental work needs to be performed to evaluate these findings.

