## Supplementary Information for "A bio-informatics approach to identify new drug targets in multidrug-resistant bacteria"

Imogen Bramhill[1]*, Aryeh Chiam[1], Dominique de Jong-Hoogland[1], Martin B. Ulmschneider[1]

[1]Department of Chemistry, King's College London, 7 Trinity Street, SE1 1DB, London, UK

### Codes/Scripts

See Github page https://github.com/imogenbramhill/bioinformatics-multidrug-resistance for methods and codes used to identify membrane proteins, filter non-homologous and essential proteins, collect protein sequences, and cluster according to sequence similarity across multiple species of bacteria.

### Clustered Proteins with 90% sequence similarity

The .clstr output file from the CD-HIT suite is available as a CSV file in the linked data repository under **Dataset D1**.

### MUSCLE Sequence Alignments

109 sequence alignments (ALN files) for each cluster, as well as SecY-Sec61α subunit alignments are available in the linked data repository under **Dataset D2**.

### Additional Tables and Figures

**Table S 1** Number of transmembrane (TM) proteins containing a signal peptide (SP) next to the total number of TM proteins for each bacteria to show proportion of total membrane proteins in each species of bacteria which were excluded from further analysis due to containing a signal peptide.

| Species of Bacteria | TM proteins containing SP | Total TM proteins |
|---|---|---|
| **A. baumannii** | 109 | 787 |
| **P. aeruginosa** | 263 | 1203 |
| **E. cloacae** | 183 | 1027 |
| **E. coli O157** | 175 | 1076 |
| **K. pneumoniae** | 213 | 1140 |
| **M. morganii** | 134 | 790 |
| **P. mirabilis** | 107 | 768 |
| **P. rettgeri** | 127 | 853 |
| **P. stuartii** | 122 | 876 |
| **S. marcescens** | 205 | 1048 |
| **E. faecium** | 87 | 642 |
| **S. aureus** | 76 | 676 |
| **H. pylori** | 45 | 318 |
| **C. jejuni** | 46 | 360 |

| | | |
|---|---|---|
| **S. typhimurium** | 146 | 1001 |
| **N. gonorrhoeae** | 71 | 383 |
| **S. pneumoniae** | 73 | 538 |
| **H. influenzae** | 42 | 346 |
| **S. flexneri** | 124 | 823 |
| **S. dysenteriae** | 133 | 871 |
| **M. tuberculosis** | 101 | 727 |

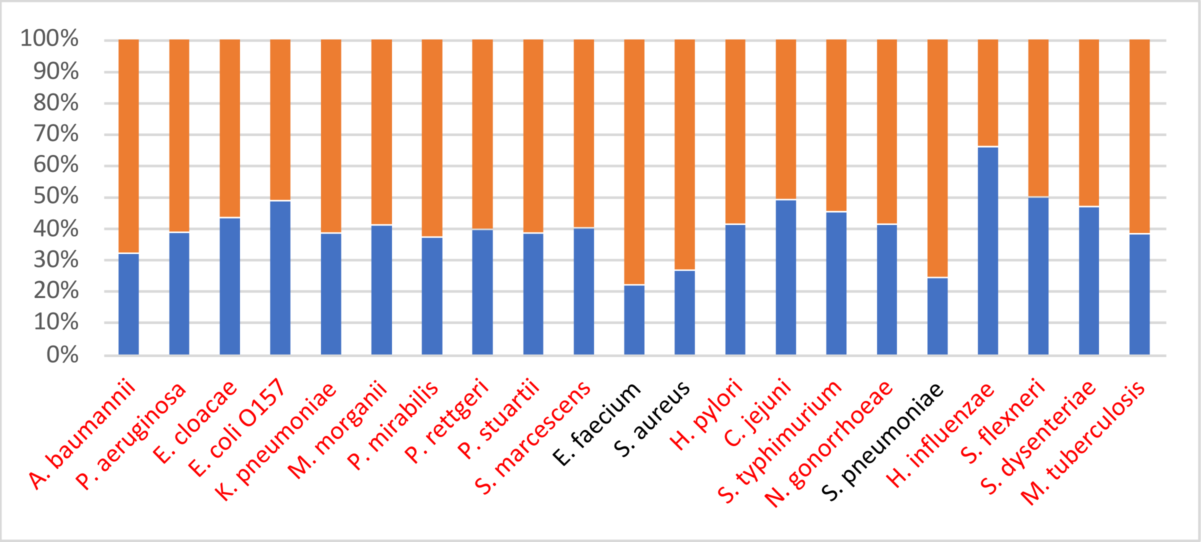


**Figure S 1** Bar chart showing the number of essential proteins, represented in blue, as a fraction of the total human non-homologous membrane proteins, non-essential proteins are represented in orange; gram-negative bacteria labels shown in red, gram-positive bacteria labels shown in black.

**Table S 2** Protein names, populations, and transmembrane topology predictions of the 109 clusters containing ≥ 5 proteins with 90% sequence similarity. Additional notes to justify excluding the potential hit from further analysis.

| Cluster Number | Protein Name (extracted from genome annotation) | No of proteins in cluster | No of Helices (majority) – taken from Phobius | Additional Notes |
|---|---|---|---|---|
| **47** | multidrug efflux system protein/pump AcrB | 6 | 12 | |
| **79** | Valyl-(-)tRNA synthetase/ligase | 6 | 1 | |
| **120** | glucosyltransferase MdoH | 6 | 8 | |
| **122** | Penicillin-binding protein 1B; bifunctional glycosyl transferase/transpeptidase | 5 | 1 | |
| **204** | pyruvate/proton symporter BtsT; carbon starvation protein (A) | 6 | 16 | |
| **230** | carbon starvation protein (A)/(CstA) | 5 | 16 | |
| **364** | PTS mannitol transporter subunit IICBA | 5 | 8 | |
| **371** | Penicillin-binding protein 2/peptidoglycan DD-transpeptidase MrdA | 6 | 1 | |
| **381** | tRNA uridine(-)5-carboxymethylaminomethyl(34) modification protein (GidA)/synthesis enzyme MnmG | 6 | 1 | |
| **465** | Division-specific transpeptidase, pencillin-binding protein 3; cell division protein/peptidoglycan glycosyltransferase/synthase FtsI | 5 | 1 | |
| **515** | PTS fructose transporter subunit IIBC | 5 | 9 | |
| **548** | (glycolate transporter); L-lactate permease | 7 | 12 | |
| **578** | ubuiquinone biosynthesis regulatory protein kinase UbiB | 5 | 2 | |
| **665** | lipid II flippase; integral m.p MviN; virulence factor; murein biosynthesis integral m.p MurJ | 6 | 13 | |
| **671** | cytochrome ubiquinol/d terminal oxidase subunit I | 6 | 9 | |
| **718** | multidrug efflux transporter MFS EmrB | 5 | 13 | |

| | | | | |
|---|---|---|---|---|
| **760** | NADH-(ubi)quinone (dehydrogenase)/oxidoreductase subunit M | 6 | 14 | |
| **793** | sodium/proline symporter PutP | 5 | 12 | |
| **799** | glycine betaine/L-proline transporter ProP | 5 | 12 | |
| **865** | lysine transporter | 6 | 12 | |
| **888** | Putative xanthine/uracil permease; adenine permease AdeP | 6 | 13 | |
| **907** | NADH dehydrogenase I subunit N; NADH-quinone oxidoreductase subunit NuoN | 6 | 14 | |
| **917** | PTS (enzyme IIsga subunit)/ascorbate transporter subunit IIC | 6 | 11 | |
| **925** | sodium/pantothenate symporter | 5 | 13 | |
| **926** | (Trk system) potassium transporter (TrkH) | 7 | 11 | |
| **973** | PTS glucose transporter subunit IIBC | 7 | 9 | |
| **1026** | D-alanine/D-serine/glycine (permease)/transporter | 6 | 12 | |
| **1080** | phenylalanine transporter | 5 | 12 | |
| **1095** | putrescine importer/symporter (PuuP)/amino acid APC transporter | 5 | 12 | PuuP found in E.coli K12 – gut bacteria |
| **1107** | (APC) amino acid transporter/permease | 5 | 12 | |
| **1148** | (phenylalanine) (APC) aromatic amino acid transporter (AroP) | 6 | 12 | |
| **1149** | Proline-specific permease ProY | 6 | 12 | |
| **1166** | (MFS) galactarate/(D)-glucarate/glycerate transporter (GudP) | 5 | 12 | |
| **1177** | putrescine importer/amino acid APC transporter | 5 | 12 | Ruled out after KEGG |
| **1208** | (sn-)glycerol-3-phosphate transporter | 7 | 12 | |
| **1232** | osmolarity sensor protein; two-component system sensor histidine kinase EnvZ | 6 | 2 | |
| **1265** | guanine/hypoxanthine permease (transporter GhxP) | 6 | 13 | |
| **1289** | (low affinity) gluconate transporter | 6 | 12 | |
| **1316** | (L-)threonine/serine transporter TdcC | 5 | 11 | |
| **1317** | preprotein translocase subunit SecY | 11 | 10 | |
| **1343** | MHS family MFS transporter | 6 | 11 | Ruled out after KEGG |
| **1394** | Lipoprotein-releasing ABC transporter LolC | 6 | 5 | |
| **1455** | HAAP family serine/threonase permease/transporter SdaC | 6 | 11 | |
| **1469** | uracil permease/transporter | 6 | 13 | |
| **1480** | C4-dicarboxylate ABC transporter DctA | 6 | 9 | |
| **1613** | Lipoprotein-releasing ABC transporter LolE | 5 | 4 | |
| **1614** | serine/threonine transporter SstT | 5 | 8 | |
| **1615** | tryptophan(-specific) transporter/permease | 5 | 11 | |
| **1716** | insertion element iso-IS10R transposase/IS4-like element ISVsa5 family transposase | 15 | 1 | protein variants in only 2 species |
| **1901** | Cell-wall shape-determining protein; rod shape-determining protein RodA; peptidoglycan glycosyltransferase MrdB | 6 | 9 | |
| **1951** | phospho-N-acetylmuramoyl-pentapeptide-transferase | 7 | 10 | |
| **1952** | lipopolysaccharide export ABC transporter permease LptG | 6 | 6 | |
| **2001** | glycine betaine/L-proline ABC transporter permease ProW | 5 | 6 | |
| **2018** | cell division protein (ABC transporter subunit) FtsX | 5 | 4 | |
| **2119** | dipeptide(/heme) ABC transporter permease DppB | 6 | 6 | |
| **2145** | (beta)-methyl-galactoside ABC transporter permease (MglC) | 5 | 8 | |
| **2156** | FtsH protease modulator/regulator HflC | 5 | 1 | |
| **2167** | fatty acid biosynthesis protein FabY; (putative) acetyltransferase | 6 | 1 | Ruled out after KEGG |
| **2222** | (high affinity) phosphate ABC transporter permease PstC | 6 | 6 | |
| **2269** | (D-)ribose ABC transporter permease | 6 | 10 | |
| **2304** | magnesium/cobalt transporter CorA | 6 | 2 | |
| **2366** | (high-affinity) branched-chain amino acid ABC transporter permease LivH | 6 | 8 | |
| **2394** | oligopeptide ABC transporter permease OppB | 7 | 6 | |
| **2436** | oligopeptide ABC transporter permease OppC | 7 | 6 | |
| **2456** | dipeptide ABC transporter permease DppC | 6 | 5 | |
| **2489** | antimicrobial peptide ABC transporter permease SapC | 5 | 6 | |
| **2491** | phosphate ABC transporter permease (subunit) PstA | 6 | 6 | |
| **2533** | sulfate/thiosulfate ABC transporter permease (subunit) CysW | 5 | 6 | |

| | | | | |
|---|---|---|---|---|
| **2574** | spermidine/putrescine ABC transporter permease PotB | 5 | 6 | |
| **2588** | PTS mannose-specific transporter subunit IID | 6 | 3 | |
| **2602** | formate transporter FocA | 6 | 6 | |
| **2643** | (spermidine)/putrescine ABC transporter permease PotI | 6 | 7 | |
| **2709** | sulfur acceptor protein CsdL; tRNA cyclic N6-threonylcarbamoyladenose(37) dehydratase/synthase TcdA | 6 | 1 | many names & 1 helix |
| **2728** | F0F1 ATP synthase subunit A | 6 | 5 | |
| **2784** | PTS mannose(/fructose/sorbose)-specific transporter subunit IIC | 5 | 7 | |
| **2802** | spermidine/putrescine ABC transporter permease PotC | 7 | 6 | |
| **2825** | (high-affinity) zinc ABC transporter permease subunit (uptake system mp) ZnuB | 5 | 9 | |
| **2836** | (lipid asymmetry maintenance) ABC transporter permease (subunit MlaE) | 6 | 6 | |
| **2944** | 1-acyl(-sn-)glycerol-3-phosphate (O-)acyltransferase | 5 | 1 | |
| **2998** | histidine(/lysine/arginine/ornithine) ABC transporter permase (HisM) | 6 | 3 | |
| **3009** | TerC family protein; transporter | 5 | 7 | Ruled out after KEGG |
| **3060** | Tol-Pal system protein/colicin transporter/uptake protein TolQ | 7 | 3 | |
| **3069** | molybdate ABC transporter permease (subunit) | 5 | 5 | |
| **3117** | glutamate/aspartate ABC transporter permease GltK | 6 | 5 | |
| **3135** | amino acid (cystine) ABC transporter permease | 6 | 3 | |
| **3150** | hypothetical protein; 7-cyano-7deazaguanine/7-aminomethyl-7-deazaguanine transporter | 5 | 6 | |
| **3162** | hypothetical protein; DedA family protein | 6 | 6 | Ruled out after KEGG |
| **3169** | glutamine ABC transporter permease GlnP | 7 | 5 | |
| **3170** | HflBKC-binding inner m.p; BAX inhibitor protein; FtsH protease modulator YccA | 5 | 7 | |
| **3202** | (DL-)methionine (ABC) transporter permease (MetI) | 6 | 5 | |
| **3301** | cytochrome o ubiquinol oxidase subunit III | 5 | 5 | |
| **3378** | Na(+)-translocating NADH-quinone reductase subunit E; electron transport complex subunit RsxA | 6 | 6 | |
| **3436** | NADH dehydrogenase subunit I; NADH-quinone oxidoreductase subunit I/NuoI | 6 | 1 | |
| **3439** | intracellular septation protein A | 6 | 5 | |
| **3502** | (pro)lipoprotein signal peptidase II | 6 | 4 | |
| **3510** | TIGR00645 family protein; hypothetical/inner membrane protein | 5 | 4 | Ruled out after KEGG |
| **3519** | (Cell wall structural complex MreBCD transmembrane component)/ rod shape-determining protein MreD | 6 | 5 | |
| **3527** | colicin V production protein | 5 | 4 | |
| **3566** | F0F1 ATP synthase subunit B | 7 | 1 | |
| **3668** | large-conductance mechanosensitive channel protein (MscL) | 5 | 2 | |
| **3686** | succinate dehydrogenase cytochrome b556 subunit | 5 | 3 | |
| **3687** | inner membrane-anchored protein; putative periplasmic protein; hypothetical protein; DUF1043 family protein | 6 | 1 | Too many unnamed proteins |
| **3732** | preprotein translocase subunit SecE | 6 | 3 | |
| **3774** | (membrane bound) cell division (leucine zipper septum) protein FtsL | 5 | 1 | |
| **3787** | fumarate reductase (membrane anchor) subunit (Frd)D | 5 | 3 | |
| **3823** | (SecYEG) preprotein translocase (auxiliary) subunit YajC | 6 | 1 | |
| **3875** | NADH dehydrogenase subunit K; NADH-(ubi)quinone oxidoreductase subunit NuoK | 6 | 3 | KEGG score < 100 |
| **3958** | F0F1 ATP synthase subunit C | 11 | 2 | |
| **3974** | S phage holin/lysis protein | 11 (all E.coli) | 2 | Ruled out after KEGG |

**Table S 3** KEGG pathway analysis of the final 109 hit proteins. Pathways are grouped by functional category, with KEGG codes and the number of mapped proteins shown.

| Category | KEGG Code | Pathway Description | Count |
|---|---|---|---|
| **Global and overview maps** | 01100 | Metabolic pathways | 18 |
| | 01110 | Biosynthesis of secondary metabolites | 3 |
| | 01120 | Microbial metabolism in diverse environments | 4 |
| | 01200 | Carbon metabolism | 2 |
| **Carbohydrate metabolism** | 00010 | Glycolysis / Gluconeogenesis | 1 |
| | 00020 | Citrate cycle (TCA cycle) | 2 |
| | 00051 | Fructose and mannose metabolism | 4 |
| | 00053 | Ascorbate and aldarate metabolism | 1 |
| | 00520 | Amino sugar and nucleotide sugar metabolism | 3 |
| | 00620 | Pyruvate metabolism | 1 |
| | 00650 | Butanoate metabolism | 2 |
| **Energy metabolism** | 00190 | Oxidative phosphorylation | 11 |
| | 00195 | Photosynthesis | 3 |
| | 00720 | Carbon fixation pathways in prokaryotes | 2 |
| | 00920 | Sulfur metabolism | 1 |
| **Lipid metabolism** | 00561 | Glycerolipid metabolism | 1 |
| | 00564 | Glycerophospholipid metabolism | 1 |
| **Glycan biosynthesis and metabolism** | 00550 | Peptidoglycan biosynthesis | 4 |
| **Genetic Information Processing** | 00970 | Aminoacyl-tRNA biosynthesis | 1 |
| | 03060 | Protein export | 4 |
| **Membrane transport** | 02010 | ABC transporters | 26 |
| | 02060 | Phosphotransferase system (PTS) | 6 |
| | 03070 | Bacterial secretion system | 3 |
| **Signal transduction** | 02020 | Two-component system | 5 |
| **Cellular community - prokaryotes** | 02024 | Quorum sensing | 6 |
| | 05111 | Biofilm formation - Vibrio cholerae | 1 |
| | 02026 | Biofilm formation - Escherichia coli | 1 |
| **Drug resistance: antimicrobial** | 01501 | beta-Lactam resistance | 5 |
| | 01502 | Vancomycin resistance | 1 |
| | 01503 | Cationic antimicrobial peptide (CAMP) resistance | 2 |

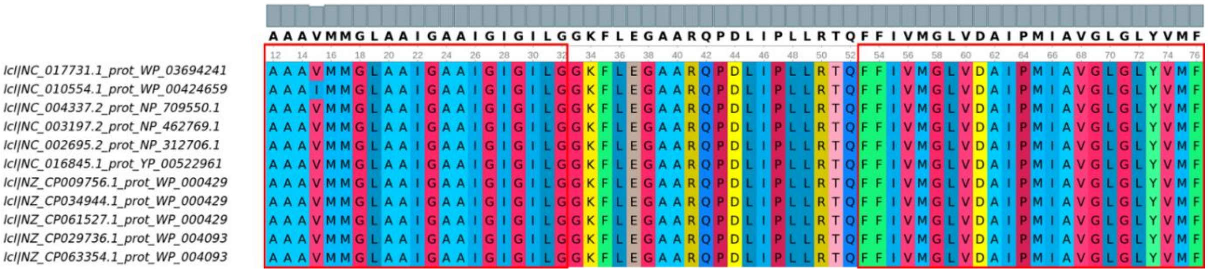


**Figure S 2** MUSCLE Alignments of F-type ATP synthase subunit C in 11 species, colour coded by residue and viewed using UGENE, with TM segments as predicted by Phobius highlighted with red boxes. The similar pattern of glycine and alanine residues in both TM segments may indicate a dimeric structure.
